# ArchiCrop: a 3D+t architectural model driven by crop model dynamics

**DOI:** 10.64898/2026.04.07.716970

**Authors:** Oriane Braud, Rémi Vezy, Thomas Arsouze, Marc Jaeger, Myriam Adam, Christophe Pradal

## Abstract

Evolving agricultural practices and contexts invite to reconsider the way crop and plant models represent agroecosystem processes. Crop models assume spatial homogeneity, which reduces confidence in their predictions for structurally heterogeneous systems, whereas FSPMs’ complexity limits their application to field-scale or large-scale studies. To benefit from strengths of both approaches, we introduce ArchiCrop, a parametric 3D architectural model of cereals that generates plant geometries constrained by crop model dynamics and coordination rules. Inspired by the concept of equifinality, ArchiCrop generates a morphospace of architecturally diverse morphotypes which remain equivalent at crop scale in terms of LAI and height. This multiscale approach enables the comparison of processes computed at different scales on wheat, rice, maize and sorghum. We demonstrate its application evaluating light interception Beer’s formalism in STICS soil-crop model relying on the leaf-resolved radiosity model Caribu for the 3D reference simulations, for a sorghum monocrop. We show that the consideration of the variability of only two plant architectural traits, leaf insertion angle and leaf number, introduces up to 27% of uncertainty in the cumulated absorbed light at the end of the season. A possible outcome from this method is also the definition of metamodels for crop model processes, as exemplified for extinction coefficient of Beer’s law. ArchiCrop can support a range of applications, including crop model uncertainty analysis, model-assisted phenotyping, and ideotype design.

**Highlights:**

- ArchiCrop is the first 3D+t botany-based parametric generative model for cereals.
- ArchiCrop downscales crop model dynamics to 3D+t architecture canopies efficiently.
- ArchiCrop compares big leaf versus leaf-resolved light interception.
- Plant architectures with same leaf area intercept light with up to 27% variability.
- ArchiCrop helps ideotyping, crop model evaluation and error propagation analysis.

**Graphical abstract:** 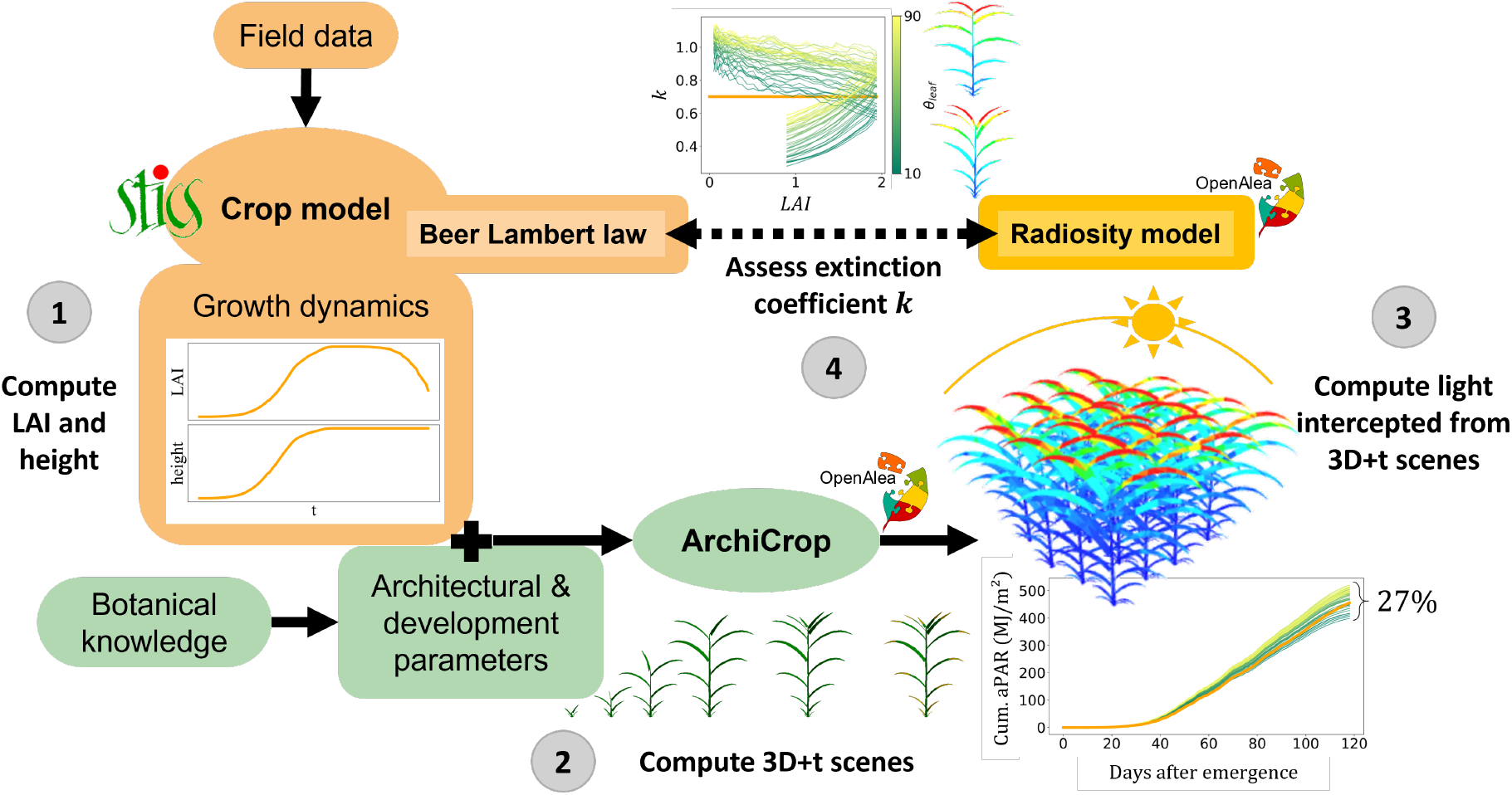

## 1. Introduction

Climate change and agroecological transition impact food production systems and their resilience in the short and long term (Bezner Kerr et al., 2023). Predictive tools are needed to understand the impact of abiotic stresses on plant production, design new agricultural system, and breed adapted new varieties (Gaudio et al., 2019; Zhang et al., 2023). Models of plant and crop growth have been widely used with these aims for decades (Gaudio et al., 2019). Yet, as agricultural systems are evolving, the relevance of the choices and assumptions made in these models has to be questioned. This is particularly relevant for diversified cropping systems that introduce spatial and temporal heterogeneity, breaking homogeneity assumptions made by crop models (Gaudio et al., 2019).

Crop models predict the impact of the environment on the whole soil-crop agrosystem by simulating key ecophysiological processes throughout the full crop lifecycle with empirical models (Boote et al., 2013; Yin et al., 2021). They consider a crop as a 1D density function and compute processes at a daily time step, which makes them computationally efficient. Crop models are calibrated on integrative field data (Pasley et al., 2022) and focus mainly on functional traits rather than structural ones (Zhang et al., 2023).

On the other hand, 3D architectural representation is explicit in structural plant models. 3D parametric architectural plant models have initially been designed in the domain of computer graphics for trees (Weber and Penn, 1995), and have been extended to annual plants, *e.g*. ADEL (Architectural model of DEveLopment based on L-systems) for maize (Fournier and Andrieu, 1999) and wheat (Fournier et al., 2003). Adding function to structure, Functional-Structural Plant Models (FSPMs) simulate 3D plant architecture from local rules to assess the role of architecture and organ-resolved biophysical processes on plant growth and function as an emergent property (Louarn and Song, 2020; Vos et al., 2007). FSPMs explicitly account for spatial heterogeneity and compute processes at a sub-daily time step, which enables to study emergent plant-plant interactions at a fine scale, but makes them computationally intensive (Gaudio et al., 2019).

Hybrid models, *i.e*. crop models that explicitly represent organ topology, have been developed to account for leaf and tiller development and carbon allocation ( EcoMeristem (Larue et al., 2019) and SiriusQuality (Martre et al., 2015)). Other models, like GreenLab (de Reffye et al., 2021) and APSIM (Brown et al., 2014), simplify organ representation as cohorts to differentiate different development stages. However, these models do not consider 3D representation of organs, neither compute plant-environment interaction in 3D, such as light interception. Individual-based models, such as FlorSys (Colbach et al., 2021), are also worth mentioning as they investigate crop heterogeneity and plant-plant interactions with simplified representations of individual plants as a volumetric, voxel-based density function at plant scale rather than an explicit surface representation at organ scale.

Currently, no model combines the consideration of the whole 3D architecture, needed to represent genotype variability and crop heterogeneity, with the predictive power of existing crop modelling platforms, designed for field conditions, easy to calibrate, and thus suited for efficiently conducting large-scale studies. Such a framework is needed to assess the impact of spatial and temporal heterogeneity on crop model processes, by identifying and quantifying the sources of structural uncertainty linked to homogeneity assumptions in a crop model (Gaudio et al., 2019).

Moreover, biophysical processes are represented with different formalisms in FSPM and crop models. The representation depends on the scale considered (organ vs canopy), the assumptions made (heterogeneity vs homogeneity of the canopy), and the representation of the canopy (3D vs 1D). It is difficult to assess the role of the process representation in the final prediction, independently of other processes. Let us illustrate it with light interception, which is a major process influencing many downstream processes, such as carbon assimilation and energy balance. FSPMs typically use 3D leaf-resolved radiation interception models such as ray tracing (Hemmerling et al., 2008) or radiosity (Chelle and Andrieu, 1998) methods, whereas crop models commonly rely on Beer’s law of light extinction (de Wit, 1964), considering one (*i.e*. big leaf) or several layers in the canopy (Gu et al., 2022). The big leaf approach relies on the assumptions that the canopy behaves like a homogeneous medium of unresolved vegetation (Ponce de León and Bailey, 2019), and keeps constant structural properties throughout crop growth, by considering an integrative constant extinction coefficient *k* (Liu et al., 2021). Evaluation of light interception formalisms at crop scale requires a reference that accounts for the effects of heterogeneity on the process by observing or simulating the process at a finer scale. Several studies have done such evaluation using digitalized plants as 3D reference (Gu et al., 2022; Liu et al., 2021). However, the heavy process of plant digitalization makes the approach difficult to apply to many genotypes. Simulating 3D plant architectures with a FSPM may help overcome this issue. However, modelling process-driven growth of a plant architecture is a tedious task by itself. Consequently, previous attempts were mostly done on a static plant architecture (Barillot et al., 2011; Ponce de León et al., 2025; Ponce de León and Bailey, 2019), but a few recent studies extended the evaluation throughout the full crop growth cycle (Li et al., 2024; Pao et al., 2021), but excluding many important processes readily available in crop models.

However, instead of upscaling from organ to crop scale, we propose a novel approach that adds the 3D dimension to crop models to benefit from their strength combined with 3D architecture. To this end, we designed ArchiCrop, a parametric architectural plant model for cereals in which plant growth is mathematically constrained by crop model dynamics. While the crop model efficiently simulates the effects of multiple stresses on crop growth throughout the crop cycle, ArchiCrop generates possible 3D architectures following crop model growth dynamics, matching its functional state and estimating the differential growth of phytomers. This framework enables the computation of processes at the organ scale in 3D to challenge its implementation in the crop model that rely on the assumption of spatial homogeneity. This multiscale approach is illustrated with a use case comparing the Beer-Lambert light interception, implemented in the STICS soil-crop model (Brisson et al., 1998), with the 3D radiosity model, implemented in OpenAlea.Caribu (Chelle and Andrieu, 1998), on cereal monocrops.

## 2. Material & Method

### 2.1. ArchiCrop Model

#### 2.1.1. Model overview

ArchiCrop is a 3D architectural plant model for cereals. The model of plant 3D architecture (*i.e*. topology and geometry) and development, that we named 3D+t, is based on knowledge about coordination and allometric rules that are well conserved among families of plants, *e.g*. cereals (Evers et al., 2005). ArchiCrop can simulate a set of 3D plant architectures whose growth follows the dynamics of Leaf Area Index (LAI) and crop height simulated by the crop model. These dynamics account for stresses computed by the crop model.

A differential approach enables to constrain local differential growth of plant organs from global crop-scale dynamics. Let’s note *C*_*crop*_ the crop-scale dimension, could it be LAI or height. The first step is to downscale crop-scale growth *dC*_*crop*_ to plant growth *dC*_*plant*_ knowing the density of plants *density*. Then, differential increments in plant height and leaf area 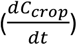 are distributed among growing phytomers *G* according to a parametrized development process, inferring organ growth *dc*_*i*_ from plant growth *dC* within a time step *dt*. ArchiCrop solves the following multi-scale equation:

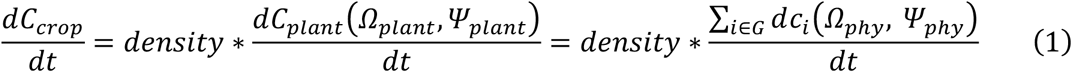

where *Ω*_*plant*_ and *Ω*_*phy*_ are respectively the sets of plant-scale and phytomer-scale parameters, and *Ψ*_*plant*_ is the set of plant-scale state variables, integrated over phytomer-scale state variables *Ψ*_*phy*_. Since the phytomer is composed of two organs, the stem and the leaf, the organ and phytomer scales are used interchangeably in this study.

Architectural and development parameters constitute degrees of freedom at the plant and organ scale with respect to the crop model. This allows to generate a morphospace of architecturally and dynamically diverse 3D+t plants following the exact same dynamics at crop scale. Our approach is inspired by the concept of equifinality, which suggests that an exact same outcome can be achieved starting from distinct initial conditions (Pasley et al., 2022).

#### 2.1.2. Cereal 3D architecture model description

ArchiCrop describes cereal plant architecture and development at organ, axis, plant, and crop scales relying on parameters. The topological representation only considers the visible part of the vegetative organs (Gauthier et al., 2020), without considering internodes, hidden sheath, and reproductive organs. We call stem element the visible part of a leaf sheath between two ligules, and leaf the leaf blade.

A Multiscale Tree Graph (MTG, (Godin and Caraglio, 1998; Pradal and Godin, 2020)) is used to represent the topology at organ, axis, and plant scales. At organ scale, a phytomer is composed of a stem element and a leaf (Figure 1A), each of which are associated with a geometric object. Each leaf branches from the top of their stem element (Figure 1A). An axis is composed (*i.e*. decomposition relation) by a sequence of phytomers (*i.e*. successor relation between stem elements) (Figure 1B). A plant is composed of a tree of axes, each axis being branched from a stem element (Figure 1C). A crop is considered as a set of plants, here positioned on a plane (Figure 1D). A schematic representation of the topology, geometry, and development at the different scales is shown in Figure 1**Erreur ! Source du renvoi introuvable**..

**Figure 1:**
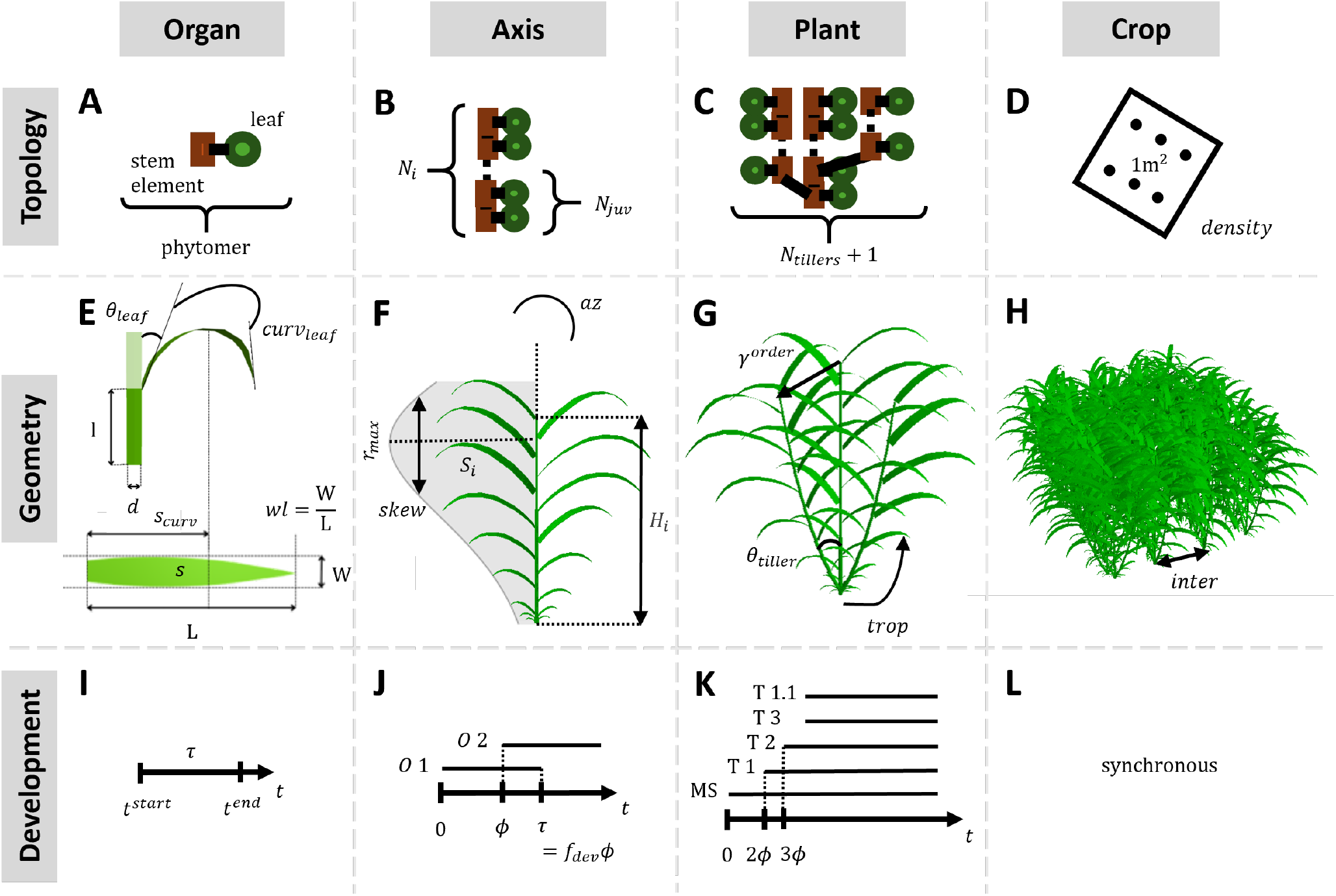
Topological (A to D) and geometrical (E to H) representations of (from left to right): an organ, an axis, a plant, and a crop, with the parameters used to describe them. Development schemes (I to L) of organs (O), axes (i.e. main stem (MS) and tillers (T)), and plants are fixed with parameters and assumed constant. Development of all plants in a crop is considered synchronous. Parameters are described in **Erreur ! Source du renvoi introuvable**..

Values for ArchiCrop architectural and developmental parameters were derived from the literature for four of the most cultivated cereal species: wheat, maize, sorghum, and rice (**Erreur ! Source du renvoi introuvable**.).

##### 2.1.2.1. Organ topology, geometry and development

###### Organ topology

A phytomer is composed of two organs, a stem element and a leaf, as well as a lateral meristem, which is implicitly considered (Figure 1A).

###### Organ geometry

A leaf is represented as a surface built by sweeping a leaf section of varying width along a midrib curve (Fournier and Pradal, 2012). The leaf width is represented using the equation of Dornbusch et al. (2011) while the leaf midrib curvature is a 2D curve defined parametrically with leaf insertion angle *θ*_*leaf*_, curvature angle *curv*_*leaf*_, and position of the inflexion point *s*_*curv*_, using the equation of Perez et al. (2016) (Figure 1E). The normalized shape is scaled for each leaf, according to a defined mature leaf area and a parameter for width-to-length ratio *wl* (Figure 1E). Leaf torsion is not considered. A stem element is represented as a cylinder described by a length *l* and diameter *d* (Figure 1E).

###### Organ development and growth

Phytomer growth and senescence are bounded in time. A phytomer starts growing at thermal time *t*^*start*^, stops growing at *t*^*end*^ after an expansion duration *τ* (Figure 1I), and starts senescing from *t*^*sen*^ after a lifespan *τ*_*sen*_. Senescence initiation depends on phenological stage, with a reduced lifespan for juvenile leaves (Brisson et al., 1998). A leaf elongates along the midrib from cell divisions at its base, so that the leaf shape appears from tip to base, where the base contains the younger tissues. During growth, leaf curvature is modelled to follow the path of the midrib curve at maturity. A leaf starts growing as erected and bends to reach given *θ*_*leaf*_ and *curv*_*leaf*_ at two thirds of leaf expansion, *i.e*. 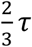, such that, at any thermal time 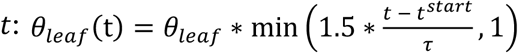, and respectively for *curv*_*leaf*_ . A stem element grows in length and in diameter, such that its diameter *d*(*t*) grows from a quarter of its mature diameter *d* to reach it after 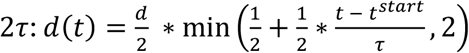.

##### 2.1.2.2. Axis topology, geometry and development

###### Axis topology

An axis is a sequence of *N*_*i*_ phytomers, ordered by their rank *r* (Figure 1B).

###### Axis geometry

The geometry of an axis is represented by a frustum decomposed into continuous elements (Pradal et al., 2009). The diameter varies along the axis linearly from base to top diameters, *d*_*base*_ and *d*_*top*_. The phyllotaxy of the leaves is defined by an azimuth angle *az* (Figure 1F).

The distribution of organ dimensions along axes follow allometric laws that model leaf and axis coordination and expansion (Andrieu et al., 2006; Martre and Dambreville, 2018). These are defined as normalized laws, discretized to the number of phytomers of the axis *N*_*i*_, and scaled according to dimensions, *i.e*. leaf area and height, for a mature axis, respectively *S*_*i*_ and *H*_*i*_.

The distribution of ligule heights is defined by parts: the *N*_*juv*_ juvenile phytomers stay small whereas later phytomers elongate (Figure 1F). The stem elements of juvenile phytomers all have the same length *l*_0_, *i.e*. the height difference between two ligules is *h*_*r*_ − *h*_*r*−1_ = *l*_0_ for *r* ∈ [1, *N*_*juv*_], such that the height of all juvenile stem elements is *H*_*juv*_ = *l*_0_ ∗ *N*_*juv*_. The heights of the ligules of later phytomers *h*_*r*_ follow a geometric distribution of common ratio *q*, such that, for later phytomer of rank *r*:

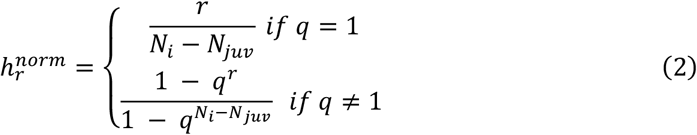

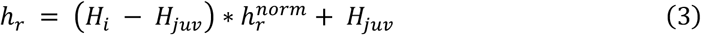

Stem elements are the parts of the sheath between ligules, so the length *l*_*r*_ of a stem element of rank *r* is determined by the difference between the *h*_*r*_ and *h*_*r*−1_.

The distribution of leaf areas along an axis follow a bell shape distribution defined with parameters *r*_*max*_ and *skew* (Figure 1F) (Fan et al., 2021). This relative leaf area distribution *s*(*r*) is discretized as a function of leaf rank *r* on an axis:

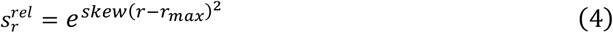

The leaf area 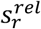of all leaves of rank *r* on axis *i* are then normalized to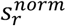, such that the sum of all leaf areas of the axis is equal to 1:

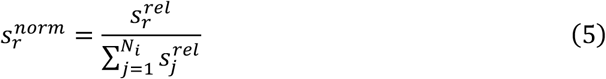

Finally, mature leaf area at rank *r* on axis *i* is obtained by scaling normalized area 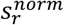 with axis leaf area *S*_*i*_, defined below in Eq. 9:

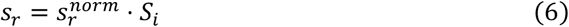

###### Axis development

Phytomers appearance occurs successively at constant thermal time interval defined by the phyllochron *ϕ* (Figure 1J). Their elongation duration *τ* can be expressed as the product of the phyllochron with a defined factor *f*_*dev*_, such that *τ* = *f*_*dev*_ ∗ *ϕ*. The period of possible growth of phytomer of rank *r* is therefore restrained to an interval 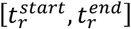, such that:

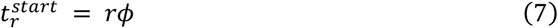

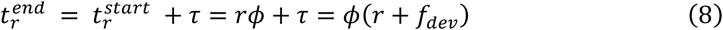

##### 2.12.3. Plant topology, geometry and development

###### Plant topology

The architecture of the plant is composed of a set of axes connected to each other as a tree structure, as described above in section 2.1.2. The first axis to grow is called the main stem. The main stem has *N*_0_ phytomers (Figure 1B). Tillering, *i.e*. basipetal ramification, can occur when lateral meristems develop into new axes. Number of tillers is set by the parameter *N*_*tillers*_ (Figure 1C). The order of an axis *i* (*order*_*i*_) is defined recursively. The main stem is of order 0, and any tiller has an order equal to that of its parent axis, plus one. The number of phytomers *N*_*i*_ of tiller *i* is computed from the absolute rank of its parent phytomer *r*_*parent*_ such that: *N*_*i*_ = *N*_0_ − *r*_*parent*_.

###### Plant geometry

The geometry of a plant is the union of the geometry of all its components, *i.e*. the geometry of a plant is the union of the geometry of all its axes, which in turn is the union of all the geometry of stem elements and leaves composing each axis. The size of phytomers of the main stem and tillers follow the same laws but discretized to different numbers of phytomers and multiplied by a reduction factor *γ* powered by the axis order (Figure 1G). Autosimilarity among tillers can be achieved by setting the reduction factor to 1 (Katayama, 1951). Plant height is equivalent to the height of the main stem. Height of tiller *i*, from its insertion, is 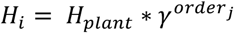. Plant-scale leaf area *S*_*plant*_ is distributed among main stem and tillers. A decreasing ratio is computed, such that the leaf area *S*_*i*_ of axis *i* of order *order*_*i*_ with *N*_*i*_ phytomers is:

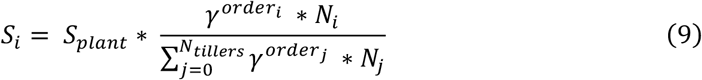

Tillers have an insertion angle of *θ*_*tiller*_ with respect to its parent axis, and bends towards the vertical according to a tropism coefficient *trop* (Figure 1G).

###### Plant development

The main stem is the first axis to start growing. Tiller emission is then defined in a deterministic way following a Fibonacci-like sequence (Goto and Hoshikawa, 1988; Toyota and Morokuma, 2021). In our model, the delay for tiller emission is constant, equal to *ϕ*, and the first tiller appears after 2*ϕ* (Figure 1K) (Evers et al., 2006). Phytomers that appear the earliest are assigned tillers, first on main stem and then recursively for phytomers of each emitted tiller, until the desired number of tillers is reached, giving priority to lower orders.

The visible part of a growing plant *Plant*(*t*) is composed of mature and growing organs *O*(*t*) (stem elements and leaves), such that, at any time *t*:

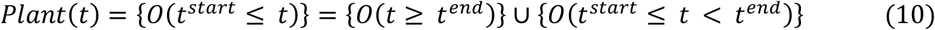

Plant leaf area *S*_*plant*_(*t*) at any time *t* is the sum of the leaf area *s*_*i,j*_ of leaves *j* of all axes *i*:

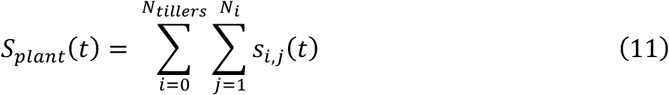

Plant height *H*_*plant*_(*t*) at any time *t* is the sum of the length *l*_*j*_ of all stem elements *j* of the main stem:

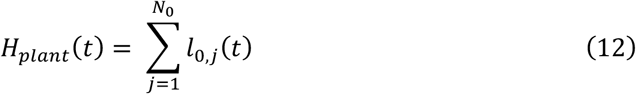

##### 2.1.2.4. Crop spatial configuration

A crop is a population of plants arranged with an explicit spatial configuration, defined by a density *density* (Figure 1D) and an inter-row distance *inter* (Figure 1H). Plant height *H*_*plant*_(*t*) is considered uniform in the canopy and equal to crop-scale height *H*(*t*). LAI is the leaf area per land surface unit. The density enables to compute the number of plants of the given species per land surface unit, such that:

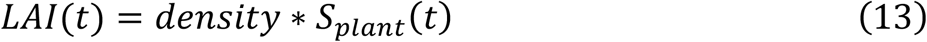

#### 2.1.3. Constraint-based growth model

In ArchiCrop, the local organ growth is constrained by crop-scale growth dynamics. Plant architecture and development are defined with parameters (Table 1), whose theoretical values for one species constitute a parameter space that we call a morphospace. However, only few combinations of parameter values or intervals enable to follow growth constraints. For parameters implied in development scheme and organ size distributions, described above in 2.1.2.1, a subspace must be defined for a given crop-scale constraint, by solving an inverse problem using the viable mathematical theory (Aubin and Frankowska, 1991), presented below. Figure 2 shows the different steps of our method to compute leaf growth of a single-stem cereal constrained by given dynamics: i) definition of a viable phytomer development scheme (Figure 2A); ii) computation of a minimal solution to follow the constrained dynamics (Figure 2B); iii) inference of viable intervals for parameters involved in organ size distributions (Figure 2C); and iv) computation of differential organ growth (Figure 2D and E).

**Table 1:**
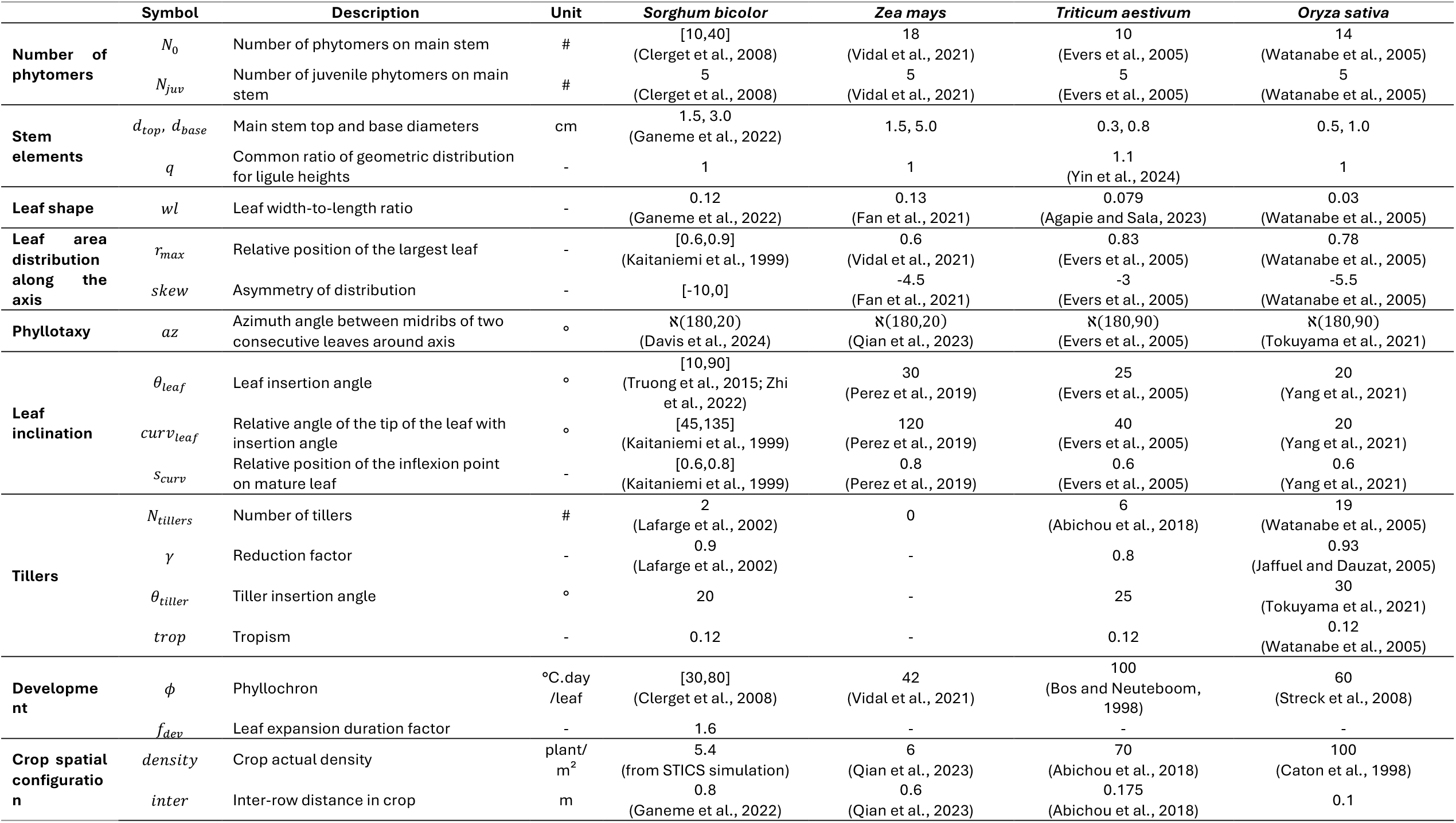
ArchiCrop architectural and development parameters used in this study, derived from literature.

**Figure 2:**
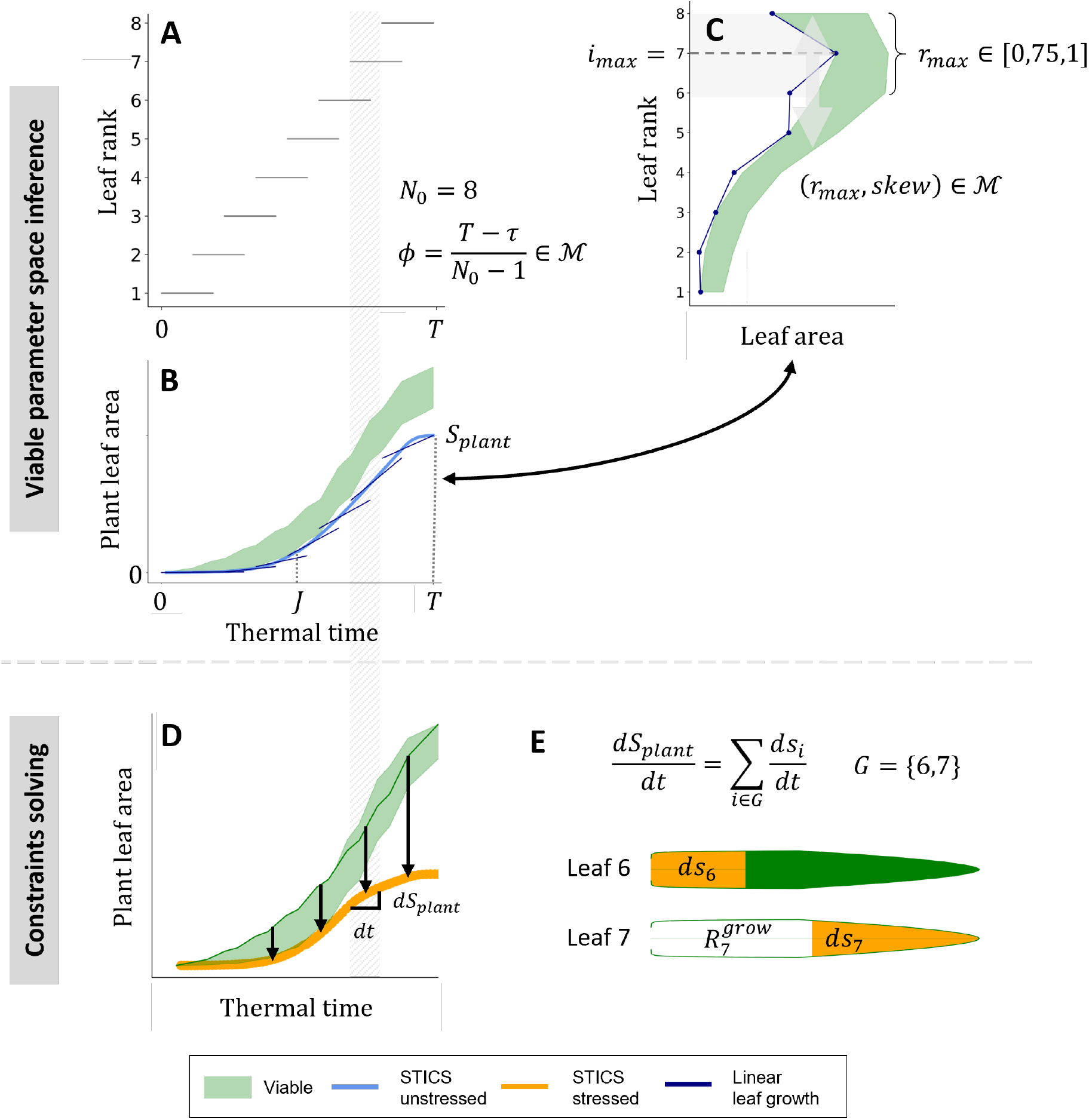
Constraint-based growth model in ArchiCrop for a single-stem cereal, showed for leaf area dynamics. **Viability parametric space inference**: A) period of phytomer development based on the phyllochron computed in the viable morphospace (ℳ); B) plant-scale leaf area growth dynamics in unstressed conditions (light blue) and minimal solution for following those dynamics assuming linear leaf growth (dark blue), and set of viable plant leaf area dynamics defined from C) assuming linear leaf growth (light green filling); C) set of viable leaf area distributions (light green filling) described by viable parameters (r_max_, skew) in ℳ (light grey filling and double arrow), above the minimal solution (dark blue). **Constraints solving**: D) reduction of viable potential growth dynamics (arrows from light green filling) to follow exactly plant-scale growth dynamics in stressed conditions (orange), by distributing differential leaf area growth 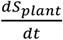) for a given dt (stripped band in D), distribution of plant-scale growth dS_plant_ (orange) among growing leaves G = {6,7}, and definition of the remaining capacity for growth of leaf 7 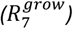 for next dt.

##### 2.1.3.1. Constraint system definition

Crop models compute growth dynamics of a crop according to a given environment and management, accounting for stresses in case of non-optimal conditions. It is often possible to deactivate the stresses for a crop model simulation, allowing to obtain a simulation at potential. In this study, unstressed simulations are therefore used to define the subspace of potential plant architectures that could grow according to the parametrization of the crop model, regardless of environmental stresses and of the ability of the crop model to consider them. Then, the growth of the potential plant architecture will be constrained to follow the growth dynamics simulated in stressed conditions at crop scale, by reducing the growth of each organ. Note that the growth of the potential plant is considered above the stressed one.

Crop models’ outputs are at crop scale. Crop-scale growth dynamics are defined by green LAI *LAI*(*t*) and crop height *H*(*t*). *LAI*(*t*) can be computed as the difference between total LAI *LAI*_*tot*_(*t*) and senescent LAI *LAI*_*sen*_(*t*) in stressed conditions. Growth dynamics of stem elements and leaves are independent in ArchiCrop: stem elements of the main stem participate to a growth in height, while leaves only account for a growth in leaf area.

Downscaling crop-scale dimension *C* implies decomposing it at plant, axis, and organ scale. The downscaling from crop scale to plant scale is explained in 2.1.2.4. Crop model dynamic outputs are given at a daily time step *dt*. In this time step, only growing organs can receive growth increments, according to Eq. 10. Growing organs of all axes are considered in the same pool *G*. Crop-scale growth dynamics are therefore distributed among plants *P* and growing organs *G* on a time step *dt* (Figure 2D-E):

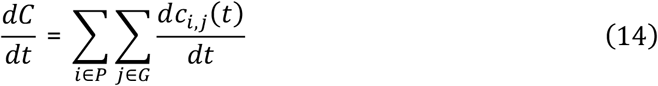

where 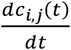 is the growth increment for organ *j* of plant *i*.

Some phenological stages of the vegetative phase can be directly determined from crop model unstressed dynamics (Figure 2B): i) emergence corresponds to the time where crop height becomes non-zero (thermal time 0 in Figure 2B); ii) the end of juvenile phase *J* corresponds to the maximal acceleration of leaf growth (Figure 2B); and iii) the end of plant growth corresponds to the time *T* when maximal LAI is reached (Figure 2B).

##### 2.1.3.2. Viability parametric space inference

The full range of parameter values for a given species defines the full theoretical morphospace of possible plant architectures. Crop-scale dynamic constraints restrict this space, defining an effective subspace. However, sampling across this reduced morphospace may yield morphotypes that fall outside this constrained subspace, due to a lack of synchronism between parametrized development and constrained growth. There is therefore a need to define a viable morphospace, named the viability kernel (Aubin and Frankowska, 1991), composed exclusively of potential plants whose architectural parameters enable to satisfy growth constraints (Figure 2B).

A set of genotypic parameters (**Erreur ! Source du renvoi introuvable**., except crop spatial configuration) defines potential plant-scale leaf area and height dynamics (Figure 2A and Figure 2C). Potential dynamics that are, at any time, greater than the unstressed dynamics are considered viable, because it is possible to obtain an exact solution by reducing the growth of organs. The aim is to find the sets of parameters that give viable dynamics (light green filling in Figure 2B), such that:

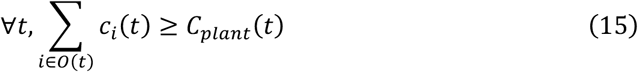

where *c*_*i*_(*t*) is the potential organ size of organ *i* at time *t, C*_*plant*_(*t*) is the plant-scale constraint at time *t*, and *O*(*t*) is the set of organs appeared at time *t* (*cf* Eq. 10), according to parametric development scheme.

Relationships between parameters enable to limit the interval of their possible values. Fixing the organ elongation duration *τ* and the number of phytomers *N*_0_, the possible value for the phyllochron is computed so that all phytomers can grow from plant emergence to end of vegetative growth *T*, based on Eq. 8, using the following relationship (Figure 2A):

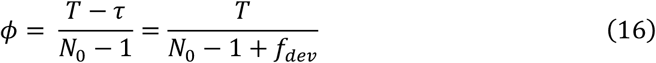

Viable values for *ϕ* have to remain within the theoretical range for a given species (**Erreur ! Source du renvoi introuvable**.).

For single-stem plants, a possible solution of Eq. 15 for a *N*_0_-phytomer plant that follows the plant-scale constrained dynamics *C*_*plant*_(*t*) can be analytically obtained by assuming a linear growth for phytomers (Figure 2B). Hence, for a phytomer of rank *r* of dimension *c*_*i*_:

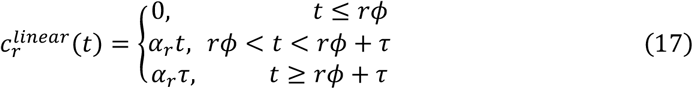

where *α*_*i*_ is the emergent growth rate of phytomer of rank *r*, and *α*_*r*_ > 1, so that, at each time *t*:

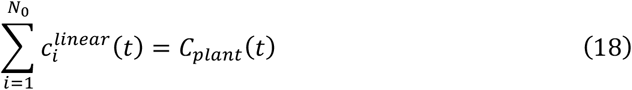

Therefore, the plant-scale dimension *C*_*plant*_ can be evaluated at times corresponding to the end of elongation of each phytomer, and the mature dimension of each organ of a plant following the constrained dynamics can be computed by resolving the following system of equations (Figure 2B):

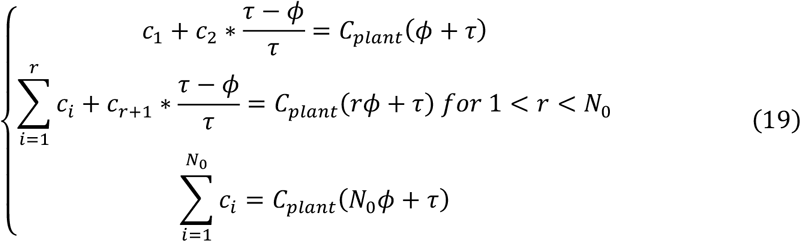

This linear system can be expressed in a matrix form *AX* = *B*, where:

- *A* is a *N*_0_ × *N*_0_ matrix carrying the advancement of elongation of phytomers *i* at the end of elongation of phytomers *j*, such that:

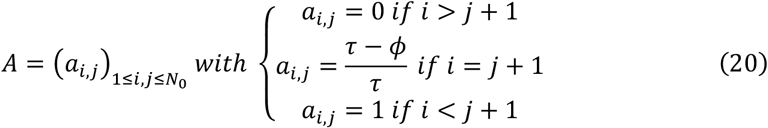
- *X* is the vector of unknown phytomer dimensions: 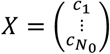
- *B* is the vector of constrained plant-scale dynamics evaluated at end of elongation of phytomers of the main stem: 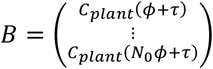

Eq. 19 for a single-stem plant is solved using the solver linalg.solve from numpy library (van der Walt et al., 2011).

When considering a cereal plant with tillers, plant height still corresponds to the height of the main stem, while the plant-scale leaf area is distributed between the main stem and all tillers, knowing their development stages, and thus, the number of growing leaves. The advancement of leaf elongation of each tiller is added to *A*, such that:

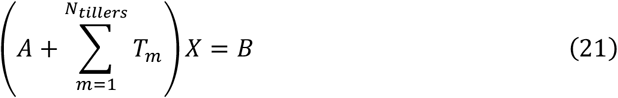

Tillers follow the same shape as the main stem, discretized in fewer phytomers, such that:

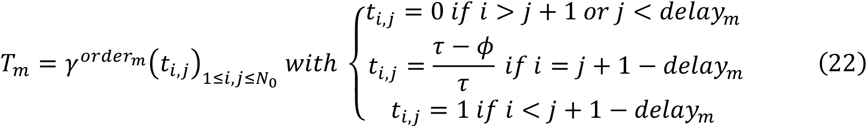

In this case, there is no analytical solution. A Non-Negative Least Squares (NNLS) solver (Lawson and Hanson, 1995) is used for Eq. 21. The solver gives the minimal solution for phytomer dimensions that follows the plant-scale constrained dynamics (leaf areas in dark blue in Figure 2B and C). In order to build a viable plant with ArchiCrop, the interval of values for the parameters defining distributions of potential organ dimensions along an axis can then be deduced.

For leaf area distribution, a plausible interval for relative rank of the largest leaf *r*_*max*_ can be defined around the rank of the largest leaf of the linear-growing plant *i*_*max*_ (Figure 2C):

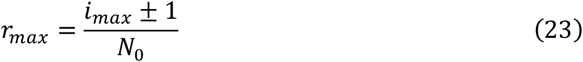

Then, for every leaf of rank *r* of the main stem, values of *skew* are computed from Eq. 4, such that:

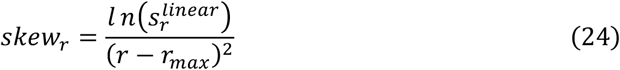

Viable pairs (*r*_*max*_, *skew*) define a potential leaf area distribution along the main stem (Figure 2C). A factor *f*_*pot*_ ensures that potential plant-scale leaf area is greater than the constraint, such that:

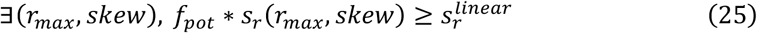

Both *r*_*max*_ and *skew* must have values that are within their theoretical range (**Erreur ! Source du renvoi introuvable**.), otherwise they are rejected.

In this section, we found the viable intervals for the phyllochron and organ size distribution parameters, that allow to simulate a viable morphospace.

##### 2.1.3.3. Constraints solving

###### Distribution of growth among growing organs in a plant

Based on phytomer development parameters, the time interval when a given organ is growing and senescing is known (Figure 2D). At each time step *t*, organs are classified according to their developmental status: not appeared, growing, mature, or senescent. Leaves and stem elements are considered growing organs if their chronological age lies between initiation 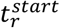 and full expansion 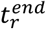 (Eq. 7 and 8), and if their visible dimension *c*_*r*_(*t*) is lower than their potential mature dimension *c*_*r*_. Leaves are senescing from the onset of senescence 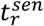, and until the organ has fully senesced. Daily growth is distributed among growing organs, and daily senescence among senescing organs (Figure 2E). The remaining capacity for expansion 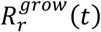 or senescence 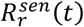 of an organ of rank *r* at a given time *t*, respectively in 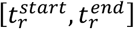 and 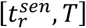, is given by: 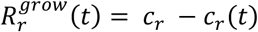 and 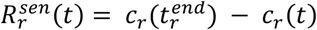 (Figure 2E).

Let’s consider a distribution function *F* of plant-scale increments among *g* growing or senescing organs, such that: 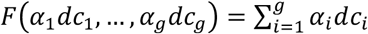, where 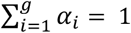 . An equal distribution would assume that all *g* organs receive an identical fraction of the increment, so 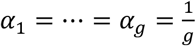. The allocation strategy used for distributing the plant-scale growth increments *dC*_*plant*_(*t*), *i.e. dS*_*plant*_(*t*) for leaf expansion, *dH*_*plant*_(*t*) for stem elongation, is a demand-based distribution. The demand-based distribution assumes that each organ receives a fraction proportional to its remaining capacity, such that 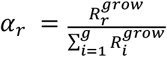. The strategy used for distributing the plant-scale leaf senescence increment *dS*_*sen,plant*_(*t*) is an age-based distribution. The age-based distribution, for senescing organs, assumes that older leaves are senescing faster than younger ones, such that 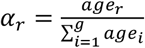.

The actual growth or senescence increment given to an organ is restrained by its remaining capacity, such that: *dc*_*r*_(*t*) = min(*α*_*r*_ ∗ *dC*(*t*), *R*_*r*_ (*t*)). If the increment for an organ is superior to its remaining capacity (*i.e. α*_*r*_ ∗ *dC*(*t*) − *R*_*r*_ (*t*) > 0), the difference is redistributed among the remaining unsaturated organs. This process is repeated iteratively until either (i) the total increment has been fully allocated, or (ii) all organs have reached their mature size *c*_*r*_. This prevents overshooting fixed mature size. Selection of viable plant architectures prevents ending up with an unallocated quantity in case (ii), that would have been lost.

###### Phytomer geometry update after growth or senescence

Organ-scale dimension increment *dc*_*r*_(*t*) is then added to previous value of dimension *c*_*r*_(*t* − 1), and organ geometry is updated (Figure 2H). Stem elements grow in height, dimension provided by the plant-scale dynamic constrain, and in width, as explained in 0. Leaves receive an increment of area, that is converted to an increment in leaf length due to its elongation. A positive increment will lead to a proximal growth (Figure 2H), while a negative increment will lead to a distal senescence.

### 2.2. Multiscale approach for evaluating Beer’s law for light interception in a crop model

Archicrop was used as part of a multiscale approach for evaluating the implementation of Beer’s law for light interception in the STICS soil-crop model (version 10.0.0, Beaudoin et al., 2023) (Figure 3). STICS computes many processes, including light interception by the canopy using Beer-Lambert’s law, and outputs growth dynamics of crop LAI and height (Figure 3A). ArchiCrop takes as inputs crop-scale growth dynamics, and architectural and developmental parameters to compute 3D+t scenes (Figure 3B). Light interception is computed on 3D scenes using Caribu radiosity model (Figure 3C). A daily extinction coefficient *k* is computed and light interception outputs from ArchiCrop coupled to Caribu is compared to a STICS simulation using a constant *k* (Figure 3D). Note that the 3D simulations are considered as the reference.

**Figure 3:**
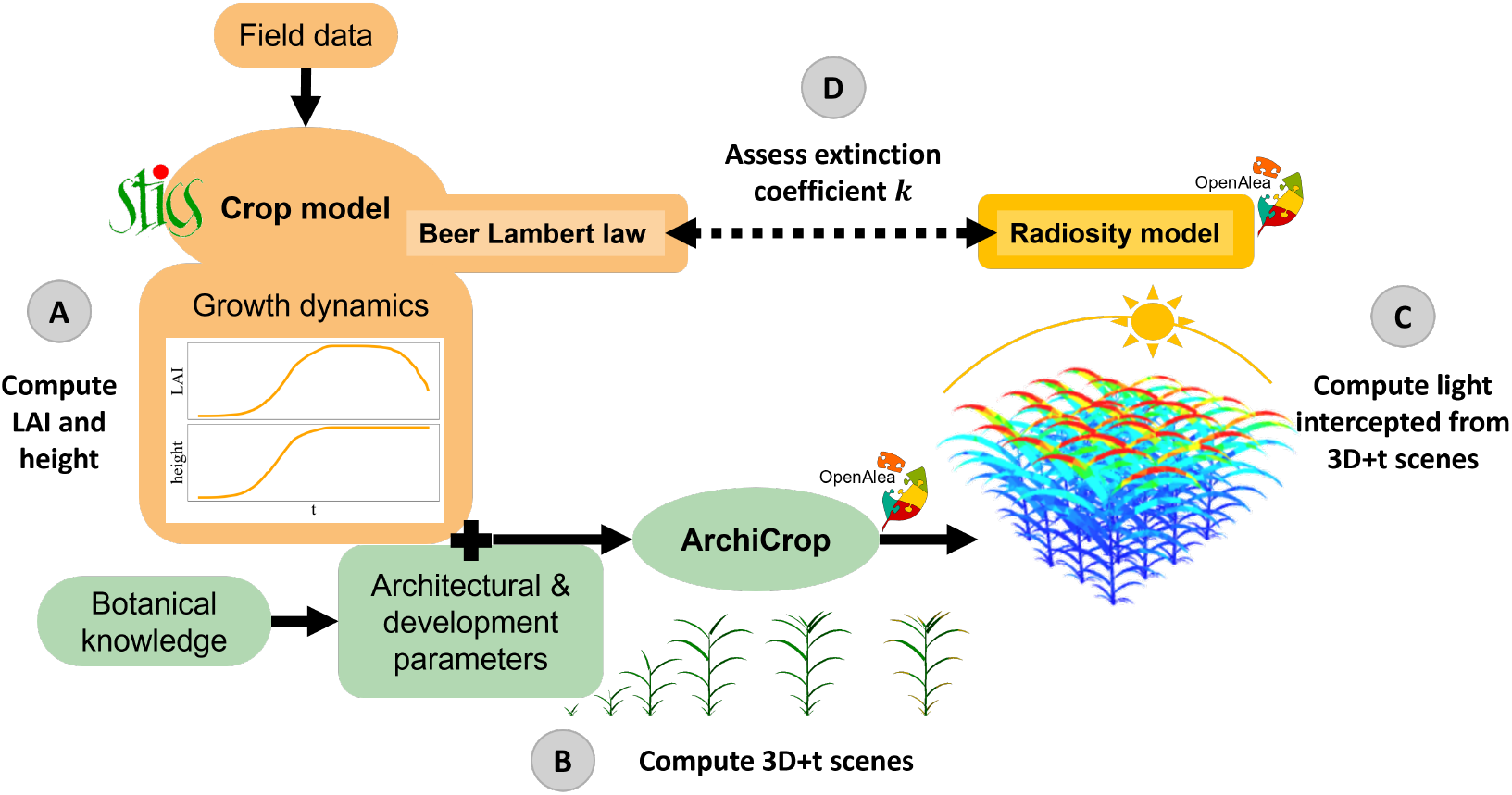
Four-step multiscale approach to evaluate light interception formalism in crop models.

#### 2.2.1. STICS simulations of *Sorghum bicolor* in Mali

A crop model simulates crop growth dynamics at a daily time step throughout the crop cycle, *e.g*. LAI (green LAI *LAI* and senescent LAI *LAI*_*sen*_, in m^2^ m^-2^), and height *H* (in m). Beer’s law computes the Photosynthetically Active Radiation absorbed by the canopy (aPAR, in MJ m^-2^ day^-1^) form the incoming PAR (iPAR, in MJ m^-2^ day^-1^):

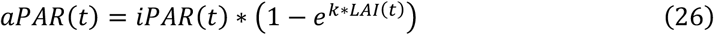

STICS is used to simulate *Sorghum bicolor* grown in N’Tarla (Sikasso Region, Mali; 12.58, –5.70) during 2018, with urea fertilization of 150 kg N/ha on Julian day 192 and 23 kg N/ha on Julian day 213 (Traore et al., 2023). Simulations are performed with and without water and nitrogen stress activation.

#### 2.2.2. 3D+t morphospace of *Sorghum bicolor* for STICS dynamics

Thanks to literature-based architectural and developmental parameters for *Sorghum bicolor* (Table 1), ArchiCrop downscales crop-scale dynamics and produces architecturally diverse 3D scenes that follow STICS dynamics, through distribution of daily crop growth among growing phytomers, as explained in section 2.1. In this study, phenotypic variability is added by varying the number of phytomers *N*_0_ between 10 and 40 and the leaf insertion angle *θ*_*leaf*_ between 10 and 90 degrees for single-stem *Sorghum bicolor* (**Erreur ! Source du renvoi introuvable**.). All other architectural and developmental parameters are fixed to values indicated in **Erreur ! Source du renvoi introuvable**., with the exception of *ϕ, r*_*max*_ and *skew* whose viable values are determined from constraints, but limited to ranges indicated in **Erreur ! Source du renvoi introuvable**.. The explicit spatial arrangement of the crop, here in-row sowing, is defined by both the effective density of plants, taken as an output of STICS, of 5.4 *plants. m*^−2^, and inter-row distance, not accounted for in STICS for monocrops, set to 0.8 *m* (**Erreur ! Source du renvoi introuvable**.).

#### 2.2.3. Leaf-resolved light interception on sorghum canopy

Light absorption by the 3D canopy, *aPAR*_3*D*_(*t*), is computed daily using leaf-resolved Caribu radiosity model (Chelle and Andrieu, 1998) implemented in the OpenAlea.Caribu library (Chelle et al., 1998). It enables to compute the light intercepted by the canopy from an incident hourly light scheme. The daily global radiation from STICS weather file is decomposed to hourly dynamics of direct light and diffuse light from a discrete hemisphere sky composed of 46 directions. Periodic boundaries are applied to the scene, effectively producing an infinite canopy.

The ratio between absorbed PAR and incoming PAR also named fraction of absorbed PAR, *faPAR*, is used for the analyses: 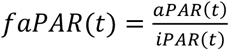.

#### 2.2.4. Quantifying the bias of Beer’s law in STICS

To evaluate the bias of Beer’s law compared to leaf-resolved 3D simulations, relative mean difference (RMD) between *faPAR* computed by STICS (*faPAR*_*Beer*_) and by ArchiCrop coupled with Caribu (*faPAR*_3*D*_) is computed:

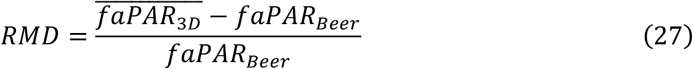

*RMD* < 1 (respectively >) means that Beer’s law in STICS underestimates (respectively overestimates) light interception by the canopy with respect to the 3D reference.

#### 2.2.5. Quantifying the uncertainty brought by plant architecture

Let *p* ∈ 𝒜 be a plant architecture *p* in the set of considered plant architectures 𝒜, 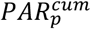 be the cumulative absorbed PAR for architecture *p*, and 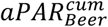be the cumulated absorbed PAR computed by the crop model using Beer’s law. Then, the architecture-induced uncertainty on cumulated absorbed PAR can be expressed as:

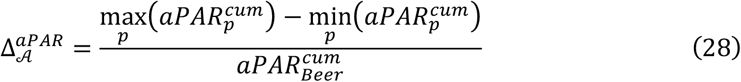

#### 2.2.6. Emergent dynamic extinction coefficient *k*

Daily values for the extinction coefficient *k*_3*D*_ (*t*) are computed from the values of *faPAR*_3*D*_ (*t*):

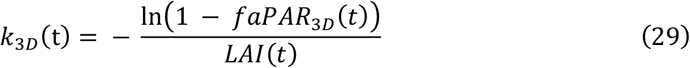

An emergent dynamic extinction coefficient can then be approximated using the following linear model:

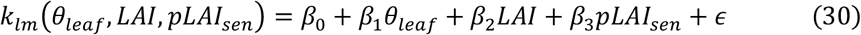

where *pLAI*_*sen*_ is the proportion of senescent LAI among total LAI, and ϵ the error.

#### 2.2.7. Model selection

The linear model for the emergent dynamic extinction coefficient computed from 3D simulations is compared to the constant extinction coefficient currently implemented in STICS. The Bayesian Information Criterion (BIC) is commonly used for model selection, penalizing higher number of parameters. Here we used it to compare Beer’s law considering a constant extinction coefficient *k*_*cst*_ and a linear extinction coefficient *k*_*lm*_, taking 3D simulations as a reference:

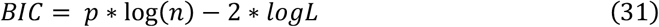

where *n* is the number of samples (*i.e*. the length of a simulations in days by the number of combinations of architectural parameters), *p* is the number of parameters (*i.e*. 1 for *k*_*cst*_ and 5 for *k*_*lm*_), and *logL* the logarithm of the likelihood function of the predicted data *faPAR*_*pred*_, such that:

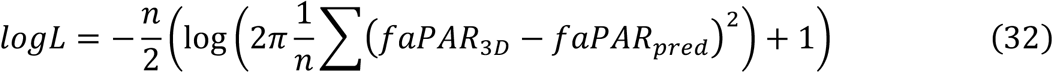

Models that lower values of BIC are generally preferred.

## 3. Results

### 3.1. ArchiCrop simulates inter-specific phenotypic variability for cereals

ArchiCrop generates the 3D architecture of the aboveground vegetative structures of four cereal species: maize, sorghum, wheat and rice (Figure 4). By varying architectural and developmental parameters within literature-based ranges for each species (**Erreur ! Source du renvoi introuvable**.), it captures interspecific phenotypic variability. We demonstrate that ArchiCrop reproduces interspecific differences from organ to canopy scale across cereals with contrasting architectures (Figure 4A).

**Figure 4:**
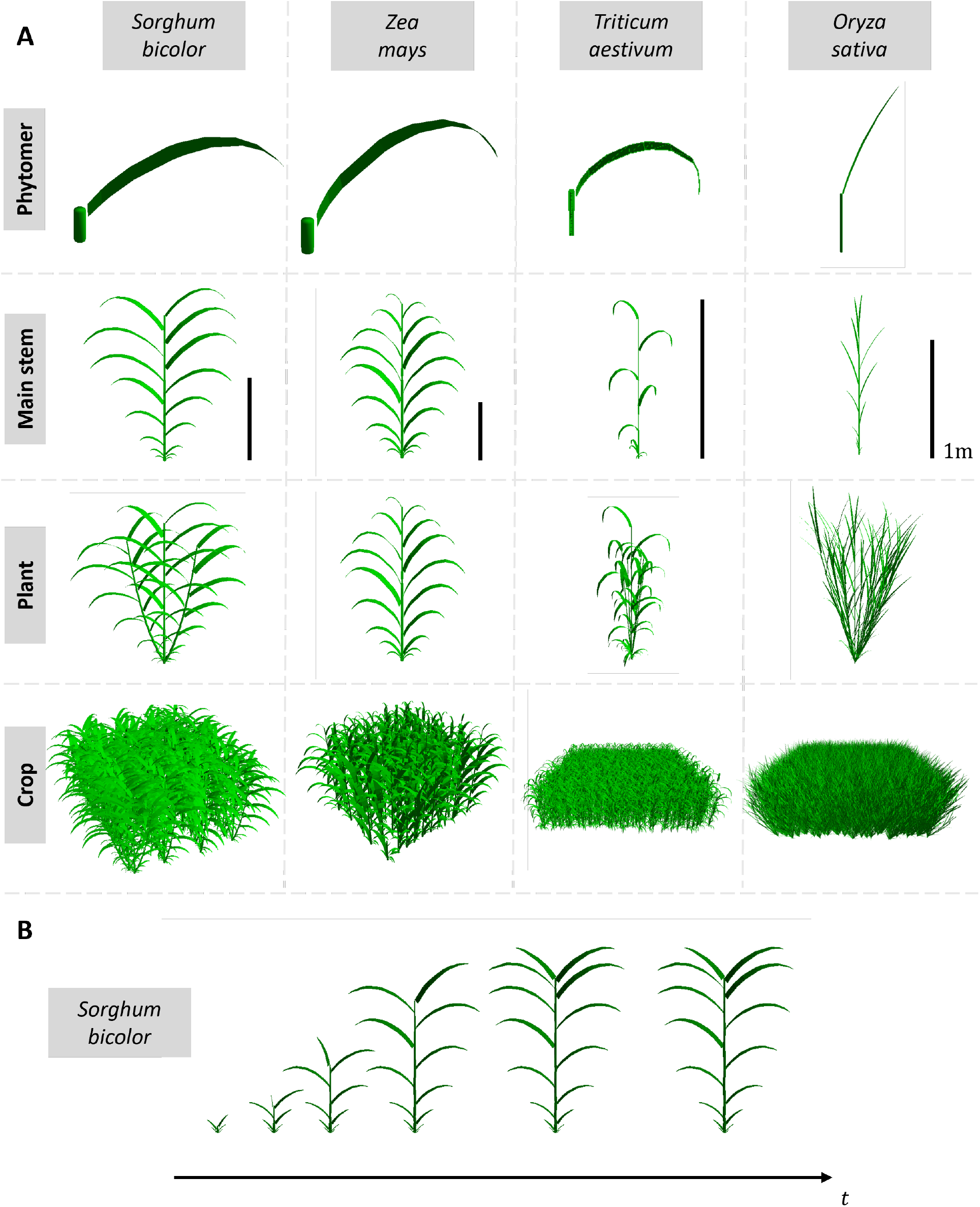
A) 3D architecture of various cereals, maize, sorghum, wheat, and rice, generated by ArchiCrop, with values in Table 1, at different scales: organ, phytomer, stem, plant, and crop. B) Example of Sorghum growth towards a mature plant following a linear growth. Differences between species arise solely from parameter values.

The architectural variability emerges across scales, from the leaf to the crop. At leaf scale, the model represents differences in length, width, slenderness, insertion angle and stiffness. At the axis scale, it reproduces distinct phyllotactic arrangements. From a limited set of traits defined at organ and axis scales, realistic plant- and crop-scale architectures emerge.

While architectural parameters define a theoretical mature plant, development parameters generate dynamics from organ to crop scale. These parameters control the timing of emergence and end of growth at all scales, together with a default linear growth pattern for each organ (Figure 4B).

### 3.2. ArchiCrop simulates 3D plants that follow crop-scale dynamics

ArchiCrop enables the exploration of the full theoretical morphospace of a species based on the combination of all ranges of parameter values, generating contrasted plant architectures. For instance, in Figure 5A, the morphospace of single-stem *Sorghum bicolor* is simulated along three dimensions by varying leaf number (*N*), phyllochron (*ϕ*), and relative position of the largest leaf along the stem (*r*_*max*_). For given crop-scale dynamics, only a subspace of the theoretical morphospace is viable (green in Figure 5A), while the remainder is not (grey), as defined in section 2.1.3.2. Viable morphotypes are those whose potential growth dynamics remain above the unstressed crop-scale dynamics (showed for LAI in Figure 5B).

**Figure 5:**
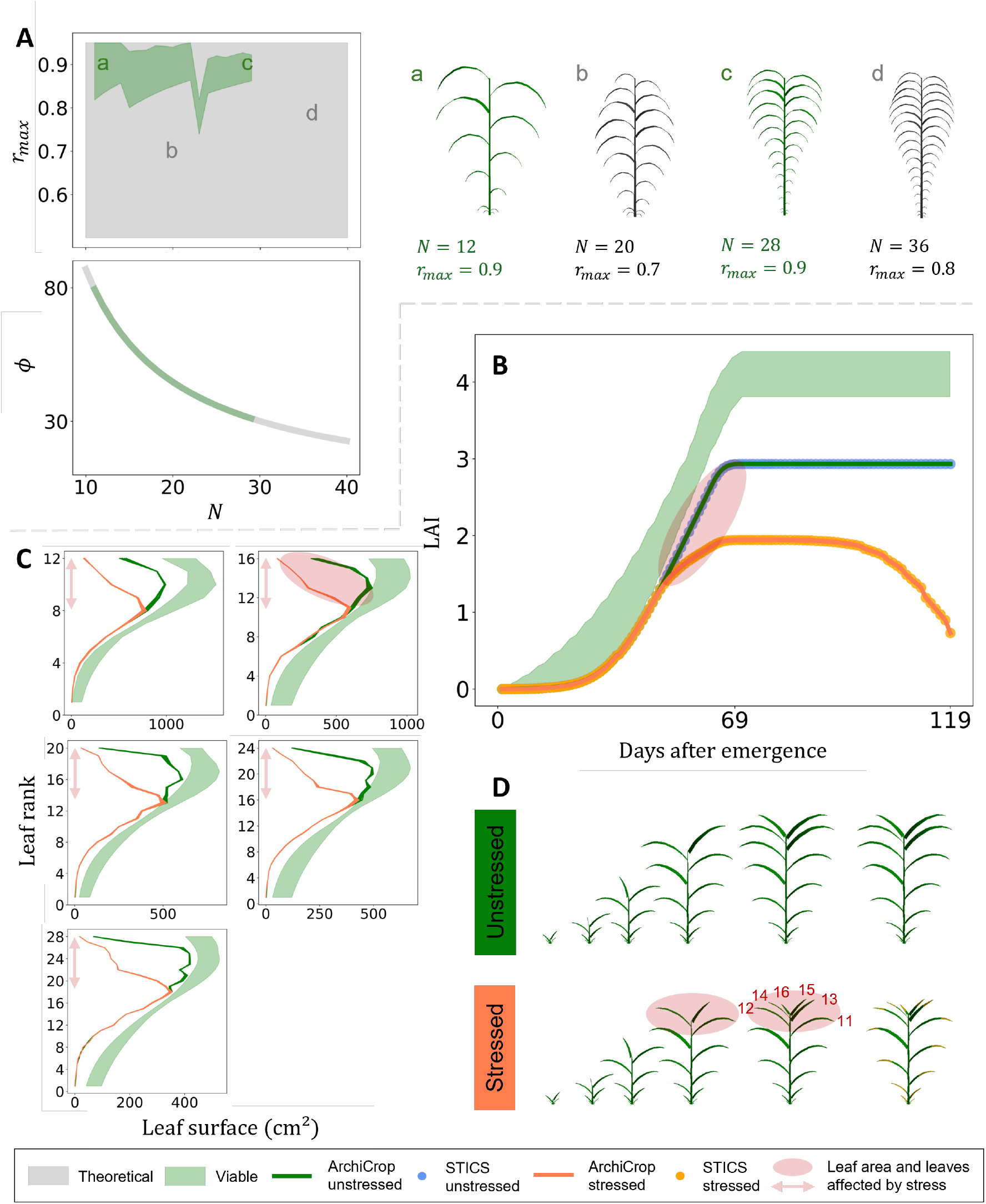
A) Three dimensions (N, r_max_, ϕ) of the theoretical morphospace of Sorghum bicolor (light grey), and the viable subspace (light green) compatible with the unstressed STICS dynamics for one site, one year. Example 3D architecture of viable (a and c) and non-viable (b and d) morphotypes are shown below. B) LAI dynamics simulated with STICS, unstressed (blue dots) and stressed (orange); potential growth envelopes of viable morphotypes (light green shading) for f_pot_ ∈ [1.0,1.5]; and growth dynamics reproduced with ArchiCrop, with (coral line) and without (green line) stress effects. C) Leaf area distributions of viable morphotypes with N = 12, 16, 20, 24 and 28, under unstressed conditions (light green shading), and simulated distributions under unstressed (green line) and stressed (coral line) conditions. The red arrows highlight the leaf ranks affected by the stress. D) 3D+t architecture of a 16-leaf sorghum plant under unstressed (top) and stressed (bottom) conditions. The light red ellipse highlights differences in leaf area under stressed conditions relative to unstressed ones.

Within this viable region, plants carry between 11 to 29 leaves, maintaining phyllochron values within a realistic range for sorghum (Figure 5A). These architectures reproduce crop-scale dynamics simulated with STICS soil-crop model under both stressed and unstressed conditions (showed for LAI in Figure 5B). Stresses simulated by the crop model and propagated to growth variables (*i.e*. LAI and height) are reflected in the emergent organ size distribution (*e.g*. leaf area in Figure 5C). Increasing the number of phytomers increases the number of leaves affected by stress. Figure 5D illustrates how these stresses translate into changes in the 3D architecture for a 16-leaf sorghum plant overtime. Note that it takes 3 seconds for simulating 119 days of organ and plant growth on a single core CPU.

### 3.3. Compare light interception formalisms at different scales of representation

Plant architecture influences light interception by the 3D sorghum canopy (Figure 6A and Figure 6B). Varying leaf insertion angle *θ*_*leaf*_ induces variations in light interception (Figure 6A). At the end of the season, the consideration of plant architecture introduced 27% uncertainty in cumulative aPAR from STICS computations (Figure 6C). Increasing *θ*_*leaf*_ leads to increasing light interception (Figure 6B) and *k*_3D_ (Figure 6D), with a slight decrease for very high values of *θ*_*leaf*_ (> 75°, Figure 6D). The variations of *k*_3D_ can be approximated by a quadratic function of *θ*_*leaf*_, 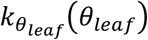, such that:

**Figure 6:**
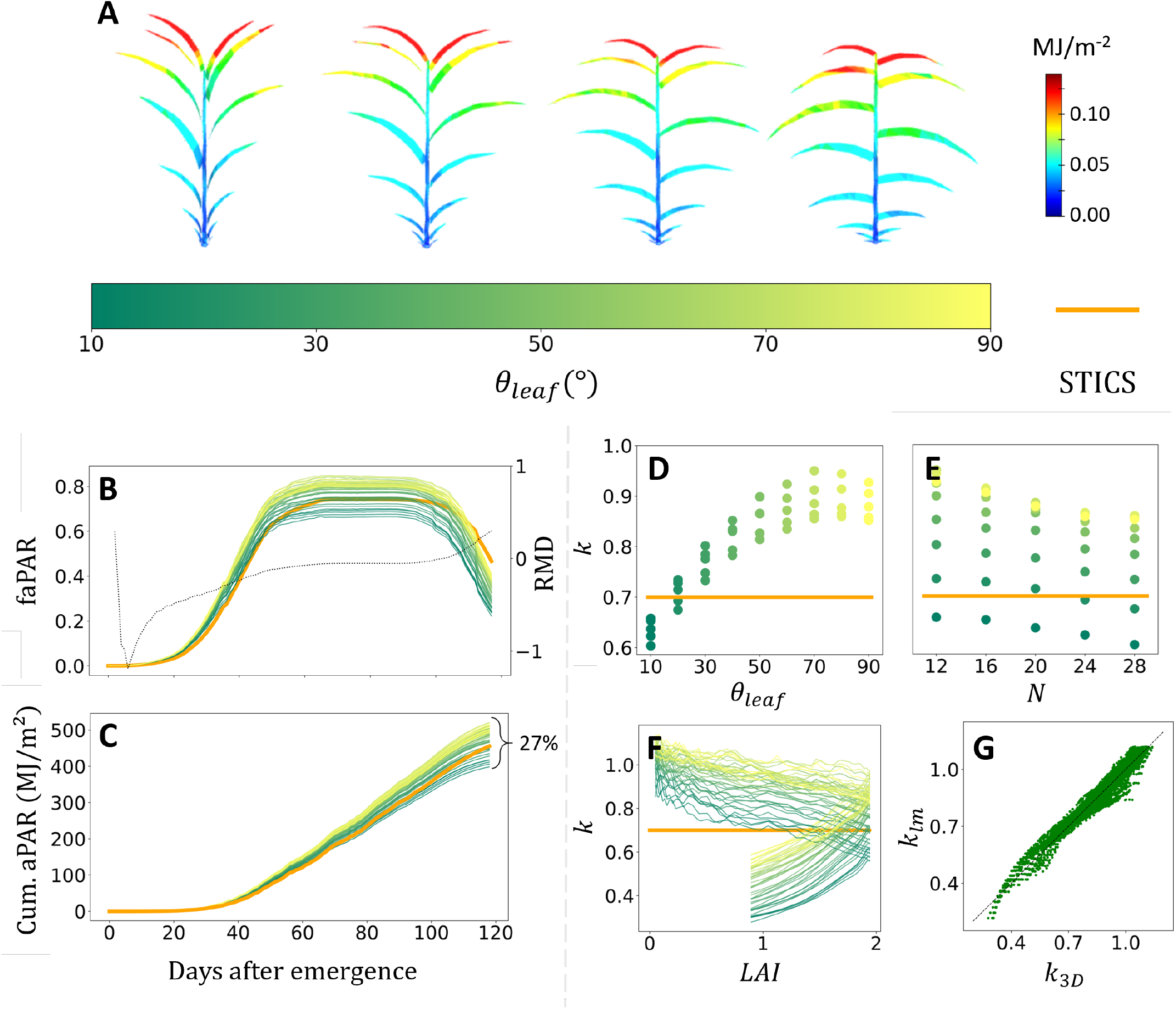
A) Daily light interception by 3D plants of 16-leaf sorghum for leaf insertion angles equal to 20, 40, 60 and 80. B) Daily faPAR computed by STICS (orange line) vs. on 3D crop for various plant architecture colored by the parameter value for θ_leaf_ (color bar). Relative Mean Difference (RMD, black dotted line) is computed throughout crop growth. C) Daily cumulated aPAR, in MJ/m^2^, computed in STICS (orange line) and in 3D for various plant architectures colored by their parameter value for θ_leaf_ (colorbar). D) Average k over the growing season as a function of θ_leaf_. E) D) Average k over the growing season as a function of N. F) Extinction coefficient: fixed value for sorghum in STICS (0.7, orange) and dynamically computed from 3D simulations for various plant architectures (colored by their value for θ_leaf_ using the color bar). We neglect the variation of extinction coefficient for very small values of LAI, i.e. for the first 20 days. G) Predicted k with linear model (k_lm_) vs. simulated k from 3D simulations (k_3D_). The black dashed line is the 1:1 line. Color bar for leaf insertion angle, in degrees, used in C, D, E, F, G.

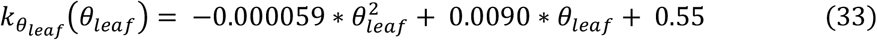

Increasing *N* leads to slightly decreasing *k*_3*D*_ (Figure 6E). The variations of *k*_3*D*_ can be approximated by a linear function of *N, k*_*N*_(*N*), such that:

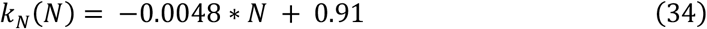

Then, doubling the number of leaves *N* in a single-stem sorghum leads to an average decrease of *k* of 0.0048, when doubling the leaf insertion angle *θ*_*leaf*_, *e.g*. from 30 to 60 degrees, increases *k* by 0.12.

STICS’s formalism is based on the Beer’s law with a constant extinction coefficient. It underestimates light interception by the canopy at the beginning of vegetative growth compared to the 3D resolution, then it converges around the average of all 3D simulations (*i.e*. RMD around 0) and eventually overestimates it during senescence phase (Figure 6B). The same phenomenon is visible when comparing fixed *k*_*cst*_ in STICS and dynamic *k*_3*D*_ emerging from 3D simulations (Figure 6F). The dynamic *k*_3*D*_ can be approximated by a piecewise linear parametric function of time *k*_*LAI*_(*LAI*(*t*)):

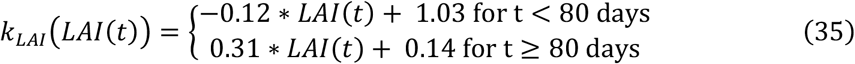

The extinction coefficient computed from 3D simulations is decreasing throughout the season (Figure 6F). For the same values of LAI, STICS does not account for distribution of green leaves in the canopy and the shading effect of senescent leaves, which has a considerable impact on *k, e.g*. for *LAI* = 1, the average *k*_3D_ is divided by 2 between vegetative growth (*k*_*LAI*_(1) = 0.91) and senescing phase (*k*_*LAI*_(1) = 0.45, Figure 6F).

Based on the results presented in Figure 6D and Figure 6F showing a large impact of green and senescent LAI along with *θ*_*leaf*_ on *k*, we fit a linear model based on these predictors to estimate a dynamic extinction coefficient *k*_*lm*_, (Figure 6G):

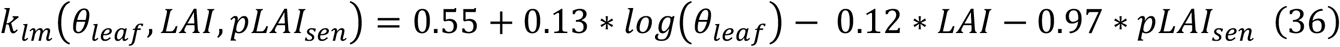

These 3 predictors can explain 94% of the extinction coefficients’ variability from 3D simulations throughout crop growth (*R*^2^ = 0.94, Figure 6G). The most influential predictors as defined by the F-statistic are (from low to high): green LAI (*F* = 1107), *θ*_*leaf*_ (*F* = 1645), and fraction of senescent LAI (*F* = 3464). *k*_*lm*_ performs better than *k*_*cst*_, with *BIC*_*lm*_ = −15661 < *BIC*_*cst*_ = −1663.

## 4. Discussion

### 4.1. ArchiCrop: a 3D+t generative model for cereals based on botanical knowledge

ArchiCrop is to our knowledge the first 3D+t parametric cereal model, able to represent various cereal species: maize, sorghum, rice, and wheat. ArchiCrop integrates the botanical structure of cereals, following both the conserved developmental pattern and the coordination rules between axes and leaves, giving it a generic nature (Evers et al., 2005). ArchiCrop can generate mockups of the most cultivated cereals, with a wide variability of architectural traits controlled by parameters (Figure 4A).

ArchiCrop can also simulate true growth, *i.e*. a plant at time *t* + 1 is relatable to a plant at time *t*. Such parametric architectural development-driven dynamic models already exist, but are often monospecific, *e.g*. (Qian et al., 2023) for maize, or adapted for two species like ADEL (Fournier et al., 2003; Fournier and Andrieu, 1999) for wheat and maize. ADEL reconstructs wheat and maize architectures by considering coordination rules and interpolating architectural constraints through time. In contrast, ArchiCrop simulates the growth of a set of 3D plants constructed from dynamic crop-scale constraints. A benefit of this hybrid approach is to simulate the function (F) by the crop model with the structure provided by ArchiCrop in an efficient way (*e.g*. 3 seconds for simulating the growth of a single-stem sorghum plant with 16 leaves, *cf* Figure 5D) using a top-down multiscale approach, rather than emerging from local laws like in FSPM. In this sense, ArchiCrop could be considered as the first hybrid FSPM or a C-FSPM (Crop-Functional Structural Plant Model), where structure is simulated according to the developmental program, and functions are simulated by a calibrated crop model.

ArchiCrop is relatively parsimonious. Only 29 parameters are required for defining the architecture and development of a species. Intraspecific genotypic variability can be represented by varying only a handful of parameter values, linked to observable changes, *e.g*. leaf insertion angle, phyllochron, number of leaves at maturity, number of tillers, parameters linked to leaf shape, distribution of the plant-scale leaf surface.

### 4.2. ArchiCrop: downscaling crop models to a dynamically constrained 3D plant model

ArchiCrop can simulate 3D plants that follow crop-scale dynamics that have been simulated by a crop model, by distributing crop-scale growth to plants in crop and plant-scale growth to growing organs in plant. Organ growth dynamics and dimensions are emergent from the simulations (Figure 5).

The parameters in ArchiCrop describing spatial arrangement, plant architecture, and development, are as many degrees of freedom regarding the crop model that does not consider them. It enables to introduce architectural and spatial variability in 3D crops that are all equivalent regarding crop-scale dynamics. Archicrop allows to explore equifinality, where different phenotypes have the same growth dynamics (Carozza et al., 2023), to understand the role of architecture on crop performance. Furthermore, the architectural variability of a given species or variety can be explored through a theoretical morphospace (Prusinkiewicz et al., 2007). This allows breeders to study the space of genotypes and traits that can produce the same plant performance, and then breed them.

A viable subspace must be identified within the theoretical morphospace to satisfy the crop-scale constraints. Biologically, this viable morphospace represents the set of all morphotypes capable of developing in accordance with the LAI and height dynamics produced by a crop model calibrated for a given variety. Defining this viable space from dynamics that exclude environmental stresses enables to find potential plant architectures consistent with the notion of “potential” commonly used in crop modelling (Pasley et al., 2022). This would allow to study in plant ecology how functional and strcutural traits are modified in different environment (Laurans et al., 2024; Violle et al., 2007).

Relationships between ArchiCrop parameters helps defining the viable morphospace. The number of leaves *N* is used to compute the phyllochron *ϕ* and parameters concerning leaf area distribution along a stem. Varying the number of leaves has therefore an effect on the distribution of leaf area among leaves in time and space (Perez et al., 2019). Relationships between ArchiCrop parameters helps defining the viable morphospace. The number of leaves *N* is used to compute the phyllochron *ϕ* and parameters concerning leaf area distribution along a stem. Varying the number of leaves has therefore an effect on the distribution of leaf area among leaves in time and space. Stresses simulated by the crop model and integrated in the growth variables (*i.e*. LAI and height) can be visible on plant architecture, mostly on the distribution of leaf area along a stem and phytomer height in ArchiCrop (Perez et al., 2019) .

### 4.3. Multiscale approach for comparing a biophysical process at crop and leaf scale

ArchiCrop can then be used in a multiscale approach to test the validity of assumptions in crop models by simulating processes at finer scale. This approach enables to consider the structural and spatial aspects explicitly, while accounting for environment and management effects on crop growth simulated efficiently by the crop model. While the crop model serves as a reference for LAI and height dynamics, emerging organ-scale simulations of processes are considered as the reference to evaluate the formalism of the given process in the crop model.

This method enables to study the added value of considering 3D features to the accuracy of formalisms of processes, as part of a crop model accounting for many processes, rather than in isolation. This allows to consider the calibration of the whole crop model and its compensations for errors (Pasley et al., 2022; Wallach, 2011). Applying this method for light interception, which is an upstream process, is a first step to analyze the propagation of error linked to plant architecture and spatial aspects in downstream processes (*e.g*. photosynthesis, C and N allocation, yield). This is often computed in studies that evaluate the effect of architectural traits or spatial features on light interception (Truong et al., 2015; Wang et al., 2026), but not as part of a whole crop model, and hence not accounting for many processes such as the effect of management or stresses.

Our approach downscales growth dynamics from crop scale to organ scale and then compares processes at these scales. However, intermediate scales and abstractions can be considered, *e.g*., for light interception, layering the canopy and computing Beer’s law on each layer (Pao et al., 2021), computing Beer’s law on voxels from an Individual-Based Model like FlorSys (Munier-Jolain et al., 2013) or using the OpenAlea.RATP model (Sinoquet et al., 2001), or considering only leaf properties in a canopy instead of a whole plant architecture (Li et al., 2024).

### 4.4. Assessing Beer’s light interception formalism in STICS crop model

We showed that the simple light interception computed with Beer’s law in STICS was in good agreement with the results from 3D simulations on varying plant architecture, but that architecture induced as much as a 27-percent uncertainty in cumulated aPAR at the end of the crop growth. These results highlight the impact of plant architecture on light interception performance, which is not accounted for in crop models, but could be with a simple linear modelling approach. This could be of particular interest for *e.g*. ideotyping, or representing the effect of stresses on leaf angle, and then model its impact on light interception.

Leaf insertion angle is known to have an important effect on light interception by a canopy (Burgess et al., 2017; Duncan et al., 1967; Perez et al., 2019; Truong et al., 2015). In our study, leaf insertion angle was a degree of freedom and was varied within a wide range for sorghum, where extreme values are less common, either because only few genotypes can display such angles (Truong et al., 2015), or because they are reached in particular environmental conditions (*e.g*. water stress, (Kenchanmane Raju et al., 2020)). However, genotypes displaying different leaf insertion angles are often sown at different densities, changing the LAI. In addition, in ArchiCrop, plant architecture is assumed not to vary with environmental conditions beyond its effects on LAI and height.

In our study, the number of leaves shows a weaker influence on light interception. Increasing leaf number leads to a slight decrease in *k*. When LAI is kept constant, a higher number of leaves implied a smaller individual leaf area, promoting deeper light penetration into the canopy, therefore explaining the reduction in *k*.

The extinction coefficient has been reported to increase with leaf angle in multiple studies (Goudriaan, 1988; Truong et al., 2015; Verhoef, 1984; Zhi et al., 2022). Unlike in some previous research (Truong et al., 2015), in this study, light interception is compared across different plant architectures at equal LAI, which makes it easier to isolate the effect of individual architectural traits. The extinction coefficient has also been shown to decrease for increasing values of LAI (Goudriaan, 1988; Li et al., 2024; Pao et al., 2021; Polley et al., 2011). Note that various values of LAI may correspond to different canopy structures or growing stages. However, during senescence, *i.e*. for a subsequent decrease of green LAI, (Li et al., 2024) reported an increase of the extinction coefficient, which could be due to not representing senescent leaves.

(Liu et al., 2021) and (Truong et al., 2015) already suggested the relevance of considering, respectively, green LAI dynamics and leaf senescence in *k*. As plants grow and senesce, the vertical distribution of green leaf area changes, and so does the light absorbed by each leaf. These dynamics are likely to play an important role during grain filling and maturation.

Although ArchiCrop represents senescent leaves, dead leaves never fall from the plant and contribute to shading until the end of the season. However, this primarily affects leaves located in the lower part of the canopy. Moreover, LAI is only distributed among leaf blades, and only light intercepted by blades is considered in our 3D simulations, though leaf sheath also intercepts light. However, LAI in crop models is calibrated with field measurements on leaf blades.

### 4.5. Perspectives

Relying on few parameters often implies considering simplifying assumptions that may impair a model’s realism. ArchiCrop considers a fixed development scheme. Studies have shown that phyllochron and leaf elongation duration could vary depending on environmental conditions, sowing dates, and leaf rank (Clerget et al., 2008). Tillering in cereals could also be complexified by considering tiller regression (Lecarpentier et al., 2019), plasticity with respect to environmental conditions, *e.g*. density and shade (Evers et al., 2006), and tiller angle dynamics (Tokuyama et al., 2021). Plasticity regarding plant height and leaf area is considered integrated in height and LAI dynamics. Moreover, the set of possible leaf area distributions within a plant, *i.e*. among axes and leaves, is explored as the viable morphospace. Other plastic deformations are nevertheless not considered in ArchiCrop but could be integrated in future versions, *e.g*. leaf insertion angle (Yang et al., 2021). Moreover, currently, the application of constraints on organs is only length-wise. However, for cereals, a stress can modify leaf width (Bouidghaghen et al., 2023), inclination and curvature (Kenchanmane Raju et al., 2020), orientation (Drouet and Moulia, 1997), rolling (Fournier and Pradal, 2012), and senescence (Robert et al., 2018). It may be interesting to extend the model to consider different deformations based on different natures of stresses, *e.g*. water, nitrogen, heat or high density, to see the role of canopy architecture on light interception as well as downstream processes.

ArchiCrop is a deterministic model and was used to represent an average plant in crop, with no intraspecific variability within the crop. Introducing intraspecific variability within a crop could be done by randomly picking architectural parameters within a distribution or adding stochasticity in the distribution of daily growth among plants in a crop, while remaining equivalent in average to the crop-scale constraint. This could enable to assess the assumptions of spatial homogeneity used within crop models.

In this study, only leaf insertion angle and number of phytomers were varied for light interception simulations. However, it has been shown that other parameters such as plant spacing (Drouet and Kiniry, 2008), may have an effect to a certain extent and with possible interactions, on variations of the extinction coefficient. Clumping factor is an integrative indicator that could be computed from 3D canopies and added as a parameter in Beer’s law (Liu et al., 2021). The linear model proposed does not have a value of truth or universality. It simply shows how we could use ArchiCrop to find a surrogate model for the extinction coefficient that could be easily integrated in a crop model, and calibrated for any cereal species using ArchiCrop. For this purpose, the surrogate model should be evaluated on other crop model simulations, *i.e*. with other crop-scale dynamics for the same species.

A long-term perspective would be to improve crop model processes with architectural parameters, without sacrificing simplicity, robustness and efficiency, while benefiting of leaf-scale models already available in OpenAlea platform, like HydroShoot (Albasha et al., 2019) and RATP (Sinoquet et al., 2001).

This hybrid strategy can be tested on various species and conditions, to hierarchize the sources of uncertainty due to architecture in the predictions of crop models, and identify their domain of validity, outside of which crop model conditions may not be fulfilled, *e.g*. in a heterogeneous crop in intercropping (Gaudio et al., 2019), or for non-linear processes like for severe drought of high temperatures in a context of climate change (Adam et al., 2020). It could be possible to analyze the error propagation due to plant architecture throughout the processes cascade of a crop model, in particular for non-linear processes *e.g*. photosynthesis, evapotranspiration, N demand. ArchiCrop can be extended to support other crop models than STICS, such as APSIM or DSSAT.

While this approach has been designed to compare plant and crop models, the exploration of a morphospace could be of great utility for breeders for ideotype design (Adam et al., 2014), *e.g*. plant traits that optimize light interception efficiency, for various cereals. ArchiCrop parameters are observable, which could ease *in silico* experiments for ideotyping. Currently, this approach has been designed for major cereals, but it could be extended to legumes, and used for intercropping systems to evaluate how crop models manage heterogeneous systems (Gaudio et al., 2019). Extending this model to other families of plants would imply knowing architectural and developmental botanical laws.

## 5. Conclusion

ArchiCrop is introduced as the first 3D+t parametric generative model for various cereal species based on botanical knowledge and crop-scale dynamics. This hybrid crop-FSPM model simulates 3D plant grow in a matter of seconds for the whole crop cycle, through a top-down approach, benefiting from the function being computed by the crop model. We introduced a multiscale approach using ArchiCrop to compare biophysical processes at crop and leaf scales. We showed a use case for light interception, comparing Beer’s law in STICS with 3D radiosity model Caribu, for a sorghum monocrop, using the 3D simulations as a reference. We found that plant architectural traits, such as leaf insertion angle, as well as leaf area dynamics and senescence have an effect on the extinction coefficient *k*.

ArchiCrop has only been studied with STICS crop model, but this approach could be applied to other crop models, or even ensemble of models. Moreover, ArchiCrop could also be used directly on phenotyping and remote sensing data to better estimate functional traits based on inversion technics (Li et al., 2025). The realism of stressed plant architectures obtained at the end of the constrained growth process could be validated using phenotyping data, comparing to 3D reconstructions, *e.g*. PhenoTrack3D (Daviet et al., 2022). Finally, ArchiCrop could be used by breeders for ideotype design by exploring the morphospace and finding plant traits that can optimize light interception efficiency for various cereals (Serouart et al., 2025).

## Code availability

ArchiCrop implementation in Python is available under the open-source license CeCILL-C on the OpenAlea platform (Pradal et al., 2008) on GitHub (https://github.com/openalea/ArchiCrop).

## Acknowledgements

This work is funded by the EU IntercropValuES project [grant number 101081973] and ANR-16-CONV-0004/#DigitAg programs. All authors have been supported by the MaCS4Plants CIRAD network, initiated by the AGAP Institute and AMAP joint research unit. We thank Christian Fournier for his advice throughout the development of ArchiCrop model, as well as his maintenance of OpenAlea Caribu model. We also thank Meije Gawinowski and Florian Larue for their advice regarding the tillering process in cereals.

## CRediT authorship contribution statement

**OB** : Writing – original draft, Writing – review & editing, Conceptualization, Formal Analysis, Investigation, Methodology, Software, Validation, Visualization. **RV** : Writing – original draft, Writing – review & editing, Conceptualization, Investigation, Methodology, Validation, Funding acquisition, Supervision. **TA** : Writing – review & editing, Conceptualization, Investigation, Methodology, Software, Validation, Supervision. **MA** : Writing – review & editing, Conceptualization, Methodology, Investigation, Validation, Supervision. **MJ** : Writing – review & editing, Conceptualization, Supervision. **CP** : Writing – original draft, Writing – review & editing, Conceptualization, Formal Analysis, Investigation, Methodology, Software, Validation, Funding acquisition, Supervision, Project administration.

## Notes

https://github.com/openalea/archicrop

### Competing Interest Statement

The authors have declared no competing interest.

